# A CyclinB2-Cas9 fusion promotes the homology-directed repair of double-strand breaks

**DOI:** 10.1101/555144

**Authors:** Manuel M. Vicente, Afonso Mendes, Margarida Cruz, José R. Vicente, Vasco M. Barreto

**Affiliations:** CEDOC - Chronic Diseases Research Center, NOVA Medical School | Faculdade de Ciências Médicas 1150-190 Lisboa, Portugal

**Author notes:** Address correspondence to: Vasco M. Barreto, CEDOC - Chronic Diseases Research Center NOVA Medical School | Faculdade de Ciências Médicas Universidade Nova de Lisboa Edifício CEDOC I (Edific. Cinzento) Rua de Câmara Pestana, n° 6 1150-190 Lisboa | Portugal.

**Keywords:** CRISPR/Cas, Gene targeting, cell-cycle, cyclinB2, HDR

## Abstract

The discovery of clustered regularly interspaced palindromic repeats (CRISPR), a defense system against viruses found in bacteria, launched a new era in gene targeting. The key feature of this technique is the guiding of the endonuclease Cas9 by single guide RNAs (sgRNA) to specific sequences, where a DNA lesion is introduced to trigger DNA repair. The CRISPR/Cas9 system may be extremely relevant for gene therapy, but the technique needs improvement to become a safe and fully effective tool. The Cas9-induced double-strand break (DSB) is repaired by one of two pathways, the error-prone Non-homologous end joining (NHEJ) or the high-fidelity Homology Direct Repair (HDR). Shifting the repair of the DSB to HDR is challenging, given the efficiency of NHEJ. Here we describe an engineered protein approach to increase knock-in efficiency by promoting the relative increase in Cas9 activity in G2, the phase of the cell cycle where HDR is more active. Cas9 was fused to the degradation domain of proteins known to be degraded in G1. The activity of two chimeric proteins, Geminin-Cas9 and CyclinB2-Cas9, is demonstrated, as well as their cell-cycle-dependent degradation. The chimeras shifted the repair of the DSBs to the HDR repair pathway compared to the commonly used Cas9. The application of cell cycle specific degradation tags could pave the way for more efficient and secure gene editing applications of the CRISPR/Cas9 system.

## INTRODUCTION

Targeted genome modifications using techniques that alter the genomic information of interest have contributed to multiple studies both in basic and applied biology. The ability to achieve localized gene targeting of the mammalian genome has provided an important research tool for probing the complex interplay between the genome, the physiologic processes it governs and the environment with which it interacts. The gene targeting procedure developed in the 1980s has been adopted by the scientific community as a routine protocol to evaluate gene function in mice [1, 2].

The finding that the generation of a double-strand break (DSB) increases gene targeting efficiency [4, 5] triggered major efforts to generate precise DSBs [5 – 8]. In response to DSBs, two cellular repair pathways are recruited to the site of the damage: the homology direct repair (HDR) pathway, where a template strand that shares homology to the damaged sequence is used in repair, and the non-homologous end joining (NHEJ) pathway, where the ligation of the two DSBs occurs in a sequence-independent manner [9 – 11]. The repair by the HDR pathway is considered to be error-free, since it entails the transfer of genetic information between the damaged DNA molecule and an intact homologous one, where the product of repair is identical to the original sequence. On the other hand, although the NHEJ pathway has the capability to ligate any given DSB ends and is considered the most efficient repair mechanism, it is error-prone due to the insertions or deletions (indels) that rise from its action.

Despite the merits of gene editing techniques invented before 2012 [6, 7, 12], the discovery of the bacterial immunity based on clustered regularly interspaced palindromic repeats (CRISPR) [13 – 15] would eventually launch a new era in gene targeting. Studies on this mechanism of defense led to the identification of CRISPR-associated genes (Cas) and the key players of the molecular response against viral genomic material [16]. The adaptive nature of the CRISPR-based immunity and how it is organized was then elucidated: an initial stage of adaptation, where the insertion of a short sequence from the viral DNA is inserted in the CRISPR locus, a second stage of CRISPR RNA (cr-RNA) maturation and finally a Cas and crRNA-mediated interference stage, during which the viral DNA is cut [17–20]. To target the DNA, it was found that the type II CRISPR system requires a single protein, Cas9, with its two nuclease domains [15, 21]. The trans-activating crRNA (tracrRNA) molecule was then found to be required for the formation of the Cas9:crRNA interference complex to direct DNA cleavage [22]. In 2012, Jinek *et al.* engineered a chimeric single-guide RNA (sgRNA), mimicking the secondary structure of the crRNA:tracrRNA, that could preserve the features required for Cas9 guidance [23], thus simplifying even more this naturally minimal system. Soon after this, three studies showed the edition of the genome of human cells, using a specific sequence to produce a Cas9-induced DSB in a targeted region of the genomic DNA [24 – 26].

HDR and NHEJ share many of the associated proteins and compete for the repair of the DSBs. One major factor influencing the pathway choice is the stage of the cell-cycle during which the DSB is introduced. Genes specific for NHEJ show constitutive expression throughout the phases of the cell cycle, leading to a constant active state for this repair pathway. In contrast, HDR is mainly active in the S and G2 stages, since a repair template becomes available, as the replication of DNA is occurring or has already occurred [27]. In S and G2, the competition between NHEJ and HDR is more accentuated, since both pathways encounter optimal conditions to act [28].

The fate of CRISPR-Cas9 induced DSBs depends on the repair pathway involved. NHEJ action promotes the formation of insertions and deletions (indels) that can be used to generate frame-shifts in the open reading frame of a given gene, often leading to loss of function (knock-out) [29 – 31]. HDR enables the introduction of any given modification at a precise location, dependent on the supply of an exogenous DNA template for repair (knock-in) [25, 32]. Producing a knock-in is more challenging than a knock-out, given the overall lower activity of HDR. Strategies to silence the NHEJ pathway have been shown to promote HDR-mediated repair, by chemically inhibiting DNA ligases, but present a rather toxic effect, due to a lack of specificity for the DNA ligase inhibited [33, 34]. Here, we describe a strategy to restrict DSB formation to the cell-cycle stages where HDR is more active, by enabling a cell-cycle-dependent degradation of Cas9.

## MATERIALS AND METHODS

### DNA and RNA extractions and cDNA generation

Genomic DNA from mouse tails or cells was extracted as described in [35]. Briefly, tail clips were incubated overnight at 50 °C in lysis buffer (50 mM Tris pH 8.0; 100 mM EDTA; 100 mM NaCl; 1% SDS), with proteinase K at 1 mg/mL. The samples were centrifuged at 16,000 rcf, the supernatant was collected and the debris was discarded. An equal volume of isopropanol (Sigma-Aldrich) was added and the solution was mixed by inverting the tubes. After a centrifugation at 16,000 rcf for 10 minutes, genomic DNA pellets were rinsed with 70% ice-cold ethanol. The tubes were centrifuged as before and pellets were left to air-dry overnight. Afterwards, they were resuspended with TE buffer (10 mM Tris pH 8.0; 0.1 mM EDTA). RNA was extracted using TRIzol (Life Technologies) and cDNA was generated using the Moloney Leukemia Virus Reverse Transcriptase (Promega), both according to the manufacturer’s protocol.

### DNA sequences and primer design

Amplification of modules was done with Phusion High-Fidelity (New England Biolabs) or Pfu Turbo polymerase (Thermo Scientific). Cyclin and Geminin modules were amplified from cDNA of the murine CH12 cell line and Cas9 from described plasmids (Addgene plasmid #43861) [36]. Primers were designed to have a region of 18-23 nucleotides of homology to a specific gene and a tail carrying overhangs with restriction enzyme sites. sgRNAs were designed with the online tool CCTop [37], ordered from Sigma, with overhangs for direct cloning in pX330 (Addgene plasmid #48137). All primers used were ordered from Sigma and all Sanger sequencing reactions were done at Stabvida.

### Cloning of murine CyclinB2 and human Geminin motifs in pX330-Cas9

A partial cDNA of human Geminin was produced from HEK293T cells using primer 5’ cagcgcctttctccgtttttctgcc 3’. The amplicon encoding amino acids 1-110 of Geminin was then amplified with primers 5’ GCGGCCGCCACCatgaatcccagtatgaagcag 3’ and 5’ CCATGGCgcttccccctcctccgcttccgccacctccgctgccccctccgccGAGCTCcagcgcctttctccgtttttctgcc 3’, adding a sequence encoding a linker. This amplicon was cloned into Addgene’s #43861 Cas9-encoding plasmid using sites NotI and NcoI (partial digest in #43861) to be expressed as a fusion to the N-terminus of Cas9, forming the 43861-Geminin-Cas9 plasmid. A partial cDNA of the murine CyclinB2 was produced from CH12F3-2 cells using primer 5’ GAGCTCgagtagcgcctccatctgcactg 3’. The amplicon encoding amino acids 2-87 of CyclinB2 was then amplified with primers 5’ GCGGCCGCCACCatggcgctgctccgacgcccg 3’ and 5’ GAGCTCgagtagcgcctccatctgcactg 3’ and replaced the Geminin encoding sequence of the 43861-Geminin-Cas9 plasmid using the sites NotI and SacI, forming plasmid 43861- CyclinB2-Cas9. Cas9 C-terminal versions with the same modules were also produced, using primers 5’ GGCCTCGAGCGCGCTGCTCCGACGCCC 3’ and 5’ GGCCTG CAGTCAGAGTAGCGCCTCCATCTGCACTG 3’ to produce the adequate CyclinB-2-encoding amplicon and primers 5’ GGCCTCGAGCAATCCCAGTATGAAG 3’ and 5’ GGCCTGCAGTCACAGCGCCTTTCTCCGTTTTTCTG 3’ to produce the adequate Geminin encoding amplicon. In both cases, the amplicons were cloned into 43861 using XhoI and PstI sites, generating 43861-Cas9-geminin and 43862-Cas9-cyclinB2.

### Cell culture and Transfections

HEK293T and U2OS cells were cultured at 37 °C, 5% CO2 in Dulbecco’s Modified Eagle Medium (DMEM; Gibco), supplemented with 10% heat-inactivated Fetal Bovine Serum (Biowest), 1 mM Sodium Pyruvate (Gibco) and Penicilin/Streptomycin (Gibco) at 100 U/mL (completed DMEM). CH12F3-2 cells were cultured at 37 °C, 5% CO2 in RPMI 1640 Glutamax medium (Gibco) supplemented with 10% heat-inactivated Fetal Bovine Serum (Biowest), 50 μM 2-mercaptoethanol, 1 mM Sodium Pyruvate (Gibco) and Penicilin/Streptomycin (Gibco) at 100 U/mL (completed RPMI).

HEK293T cells were transfected using the calcium phosphate technique. In short, cells were seeded at a density of 0.4 million cells per well in a 6-well plate and incubated as indicated, overnight. In the following day, two mixtures were prepared, in two separate Eppendorf tubes: Mix 1 – ddH2O up to 100 μL, 25.6 μL of 2M CaCl2 and 8 μg of total DNA, added in this order; Mix 2 – 100 μL of 2x Hepes buffered saline (HBS). After 1 minute of continuous air-bubbling of the HBS solution, Mix 1 was added to Mix 2 drop-wise, while flickering the tube. The mixture was left to precipitate for at least 30 minutes and then added dropwise to the cell-containing wells. Eight hours post-transfection, the media was aspirated and replaced with 2 mL of completed DMEM, pre-warmed at 37 °C.

For the cell cycle arrest, 5×10^5^ cells cells were seeded in wells of a 6-well plate and transfected in the day after, using the calcium phosphate technique previously described. 36 hours post-transfection, nocodazole was added at 100 ng/mL and the treated cells were incubated for 24 hours. Cell cycle synchronization was verified by flow cytometry, with a PI staining.

### Western Blot

G2-arrested cells were detached by mitotic shake off, and the wells were washed with prewarmed PBS, that was also collected. The unsynchronized cells were detached by trypsin incubation for 5 minutes at 37 °C. Completed DMEM was added to inactivate trypsin and cells were collected into 15 mL tubes (VWR). The tubes were centrifuged at 300 rcf for 5 minutes to pellet cells and total protein was extracted by resuspending each pellet in 1 mL of RIPA buffer (150 mM NaCl, 1% NP-40, 0.5% sodium deoxycholate, 0.1% SDS, 50 mM Tris-HCl pH 8) supplemented with protease and phosphatase inhibitor cocktail (Roche Molecular Biochemicals), and at 4 °C with gentle shaking/tumbling for 30 minutes. Tubes were centrifuged at 16,000 rcf for 10 minutes and supernatants were transferred to new tubes. Samples were stored at −20°C until analysis. Protein concentrations of the extracts was determined by the Bradford assay (Bradford reagent supplier: Bio-Rad) and the amount of total protein loaded from each sample was optimized to achieve similar GFP levels. Sample volumes were levelled with PBS (1x) up to a final volume of 30 μL and mixed with 10 μL of 4x Laemmli Buffer (50 mM Tris-HCl pH 6.8, 8% SDS, 40% glycerol, 8% 2-mercaptoethanol, 0.02% bromophenol blue) and boiled at 95 °C for 5 minutes before resolution in a 10% polyacrylamide gel, with a 6% stacking SDS gel, at 100 V for around 2 hours. Proteins were “wet” transferred to a PVDF membrane, with a pore size of 0.45 μm (Immobilon, Millipore), at 100 V for 90 minutes. Gel electrophoresis was done in a Mini-Protean Tetra system (Bio-Rad), with the corresponding module for transfer. The membranes were blocked for 1 hour with 5% milk/TBS-t, followed by overnight incubation with primary antibodies against FLAG (DYKDDDDK epitope, # AB0085-200, SICGEN) and GFP (anti-GFP, #AB0066-200,SICGEN), both at 3 μg/mL, in 5% milk/TBS-t. Membranes were incubated with HRP-conjugated secondary antibodies for both FLAG and GFP stainings (Rabbit anti-goat-HRP conjugate, #1721034, Bio-Rad) diluted at 1:5000, in 5% milk/TBS-t, and detected by enhanced chemiluminescence, with Amersham™ ECL™ (GE Healthcare), using ChemiDoc XRS System (Biorad).

### Flow cytometry

Cells were collected, centrifuged at 300 rcf for 5 minutes and pellets were washed twice with PBS containing 1% FBS. Cells were stained with antibodies at the adequate dilution for at least 20 minutes at 4 °C or on ice. After the staining, cells were washed twice and analyzed either in a FACS Aria II or Canto II (BD Biosciences, USA). Cells were interrogated: for mCherry expression using a 561 nm laser and fluorescence was measured by a 610/20 bandpass filter, and for GFP using a 488 nm laser and fluorescence was measured by a 530/30 band-pass filter, both in FACS Aria II; for PI using a 488 nm laser and fluorescence was measured by a 585/42 band-pass filter, in FACS Canto II. Flow cytometry data were analyzed using the FlowJo v.8.7 software (Tree Satr Inc, USA). The Watson pragmatic fitting algorithm of FlowJo v10.1 software was used for cell cycle analysis.

### Statistical analysis

Percentages were transformed to arcsin values and two-tailed Student’s t-tests were done to compare the means. GraphPad Prism 6 software was used to perform the analysis.

## RESULTS

### Testing Cas9:cell-cycle tags chimeric proteins

In order to restrict the activity of Cas9, i.e., the generation of DSBs, to the cell-cycle stages where HDR machinery is the most active (S and G2), chimeric proteins were engineered fusing Cas9 to the degradation domains of the human Geminin and murine CyclinB2, either to the N- or the C-terminus of Cas9 (Fig. 1A). These domains were already shown to determine a cell-cycle specific profile of chimeric proteins [38, 39], namely an increase in their relative concentration in S and G2 compared to G1, high jacking the conventional CyclinB2 and Geminin degradation pathways. The chimeric proteins were evaluated using cell lines encoding reporters in the genome that measure the efficiency of NHEJ or HR to restore the open reading frame of a GFP gene after the introduction of a DSB [40]. The DR-GFP and EJ5-GFP reporter cell lines used to assess the frequencies of repair of the Cas9-induced DSBs enabled the determination of choice of repair: HDR and NHEJ pathways. The DR-GFP reporter contains two GFP cassettes separated by 3.7 kb (Fig. 1B). The one adjacent to the promoter contains a I-SceI restriction site that gives rise to two in-frame premature stop codons, and the distal one has no mutations. A sgRNA was used to guide Cas9 to the I-SceI restriction site. If the DSB is repaired by HDR, using the intact GFP cassette, cells become GFP+. Thus, the relative numbers of GFP+ cells are a measure of HDR. In contrast, the EJ5-GFP reporter contains two I-SceI restriction sites in the extremities of a puromycin resistance (puro) gene (0.6 kb) that separates a promoter and a GFP cassette, preventing its expression. To induce DSBs in the extremities of the puro gene, sgRNAs targeting the two I-SceI restriction sites were used. If the two resulting DSBs are recombined by NHEJ and the puro gene is excised, then the cell acquires green fluorescence. C-terminal fusions were shown to be inactive in the used reporters, as indicated by the lack of ability to induce GFP-positive events (Fig. 1C). Since there is a background of GFP-positive cells (spontaneous events), the difference between the percentage of GFP+ cells when cells were transfected with Cas9 alone, or combined with the sgRNA, was determined and confirmed the previous result. In turn, the N-terminal fusions were active and were further analyzed.

**Figure 1:**
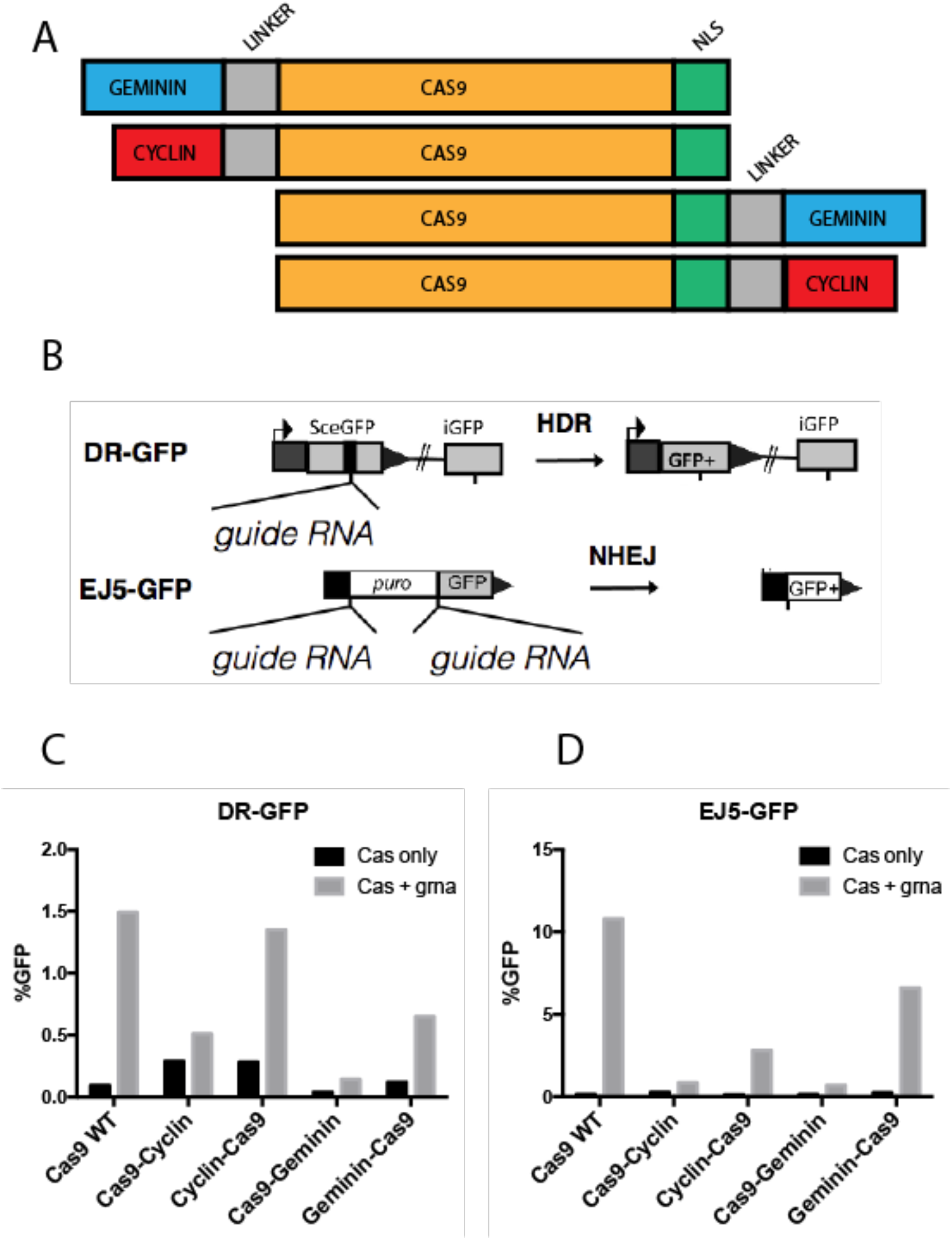
Evaluation of Cas9 fusions. (A) Scheme of the fusions tested. (B) GFP reporters for HDR, DR-GFP, and NHEJ, EJ5-GFP (C and D) Activity of fusions in the described reporters. In the DR-GFP cell line, repair of a DSB induced by Cas9 in an expressed non-functional GFP gene is done by HDR using a neighbor intact GFP gene as the repair template, giving rise to a GFP+ cell. In the EJ5-GFP cell line, the repair of two Cas9-induced DSB is done by NHEJ, which removes a piece of DNA between the promoter and GFP, allowing its expression.

### Evaluation of HDR and NHEJ activities on the repair of functional Cas9:cell-cycle tags chimeric proteins

Functional N-terminal fusions were tested for their ability to induce NHEJ and HDR events in experiments independent from the initial screen. GFP+ cells were quantified by flow cytometry and the activity of chimeric Cas9 proteins was determined for both repair processes. In the HDR reporter, all versions of Cas9, *i.e.* the wild type (WT) single Cas9 and the chimeras, drove repair events and showed similar levels of activity, with a slight increase in the case of the Geminin-Cas9 fusion (Fig. 2A). For the NHEJ reporter there were clear differences in the efficiency to generate GFP+ events. Both chimeric proteins showed a decrease in activity, when compared to WT Cas9 (Fig. 2B). Also for this reporter, the Geminin-Cas9 fusion showed higher levels of activity compared to the CyclinB2-Cas9 chimera. In G1, the chimeric Cas9 proteins are degraded and DSBs are not induced. This explains the drop in NHEJ efficiencies of the fusions compared to the WT form. Since the HDR machinery is mainly active in the S and G2 stages, and chimeric proteins are degraded in G1, the frequency of DSB induced during those stages is supposed to be the same between WT and chimeric Cas9. Thus, a useful readout is the ratio of the of HDR and NHEJ efficiencies measured by the used reporters. Chimeric proteins show an increased ratio of HDR, compared to the WT Cas9. We conclude that, compared to WT Cas9, the chimeric proteins favor repair by HDR over repair by NHEJ (Fig. 2C).

**Figure 2:**
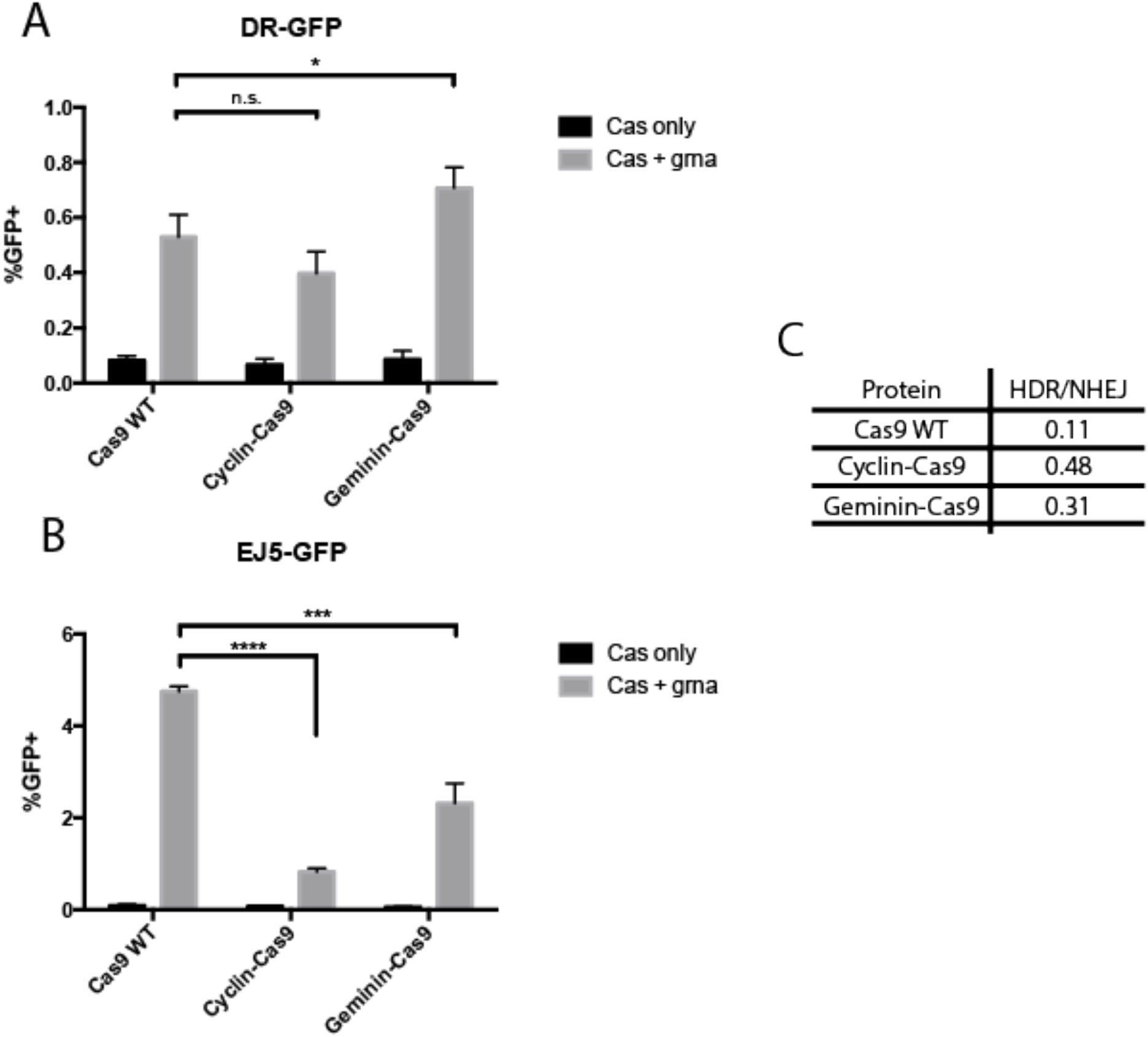
DNA repair activity on DSBs induced by chimeric Cas9 proteins. (A and B) Activity of fusions in the described reporters. The error bars show SD (n = 3). (C) HDR/NHEJ ratios for WT and chimeric Cas9. Data are representative of 2 experiments.

### Cell-cycle specific degradation of Cas9: cell-cycle tags chimeric proteins

The results of the previous section showed a bias towards HDR-mediated repair when the chimeras were used. To evaluate if this is occurring due to the predicted cell-cycle-dependent degradation of chimeras, the cells were G2-synchronized using nocodazole, a drug that prevents entry in mitosis. The drug efficiently arrested cell-cycle progression in the 293T cells used in this report, as the G2 population became almost absolute (Fig. 3A-B) Protein extracts from unsynchronized and synchronized cells were then analyzed by Western Blot. In unsynchronized cells, the levels of the chimeric Cas9 should be inferior to those of the WT, which was confirmed. Because the chimeras share the same promoter in their expression vector, this result reflects the differential proteolytic degradation of chimeras compared to the WT form or that they differ in protein folding and stability (this latter hypothesis was not further pursued). In G2-arrested cells, the degradation of chimeric Cas9 proteins should be prevented and the degradation of WT Cas9 should not be influenced. Indeed, under these conditions there is an increased accumulation of the Cyclin-Cas9 fusion, compared to the WT Cas9, normalized by the GFP loading control (Fig. 3C-D). The increase of the Geminin fusion is not so pronounced, which could explain the poorer performance of this chimera compared to the CyclinB2 chimera in favoring HDR-mediated editing events when using the GFP reporters.

**Figure 3:**
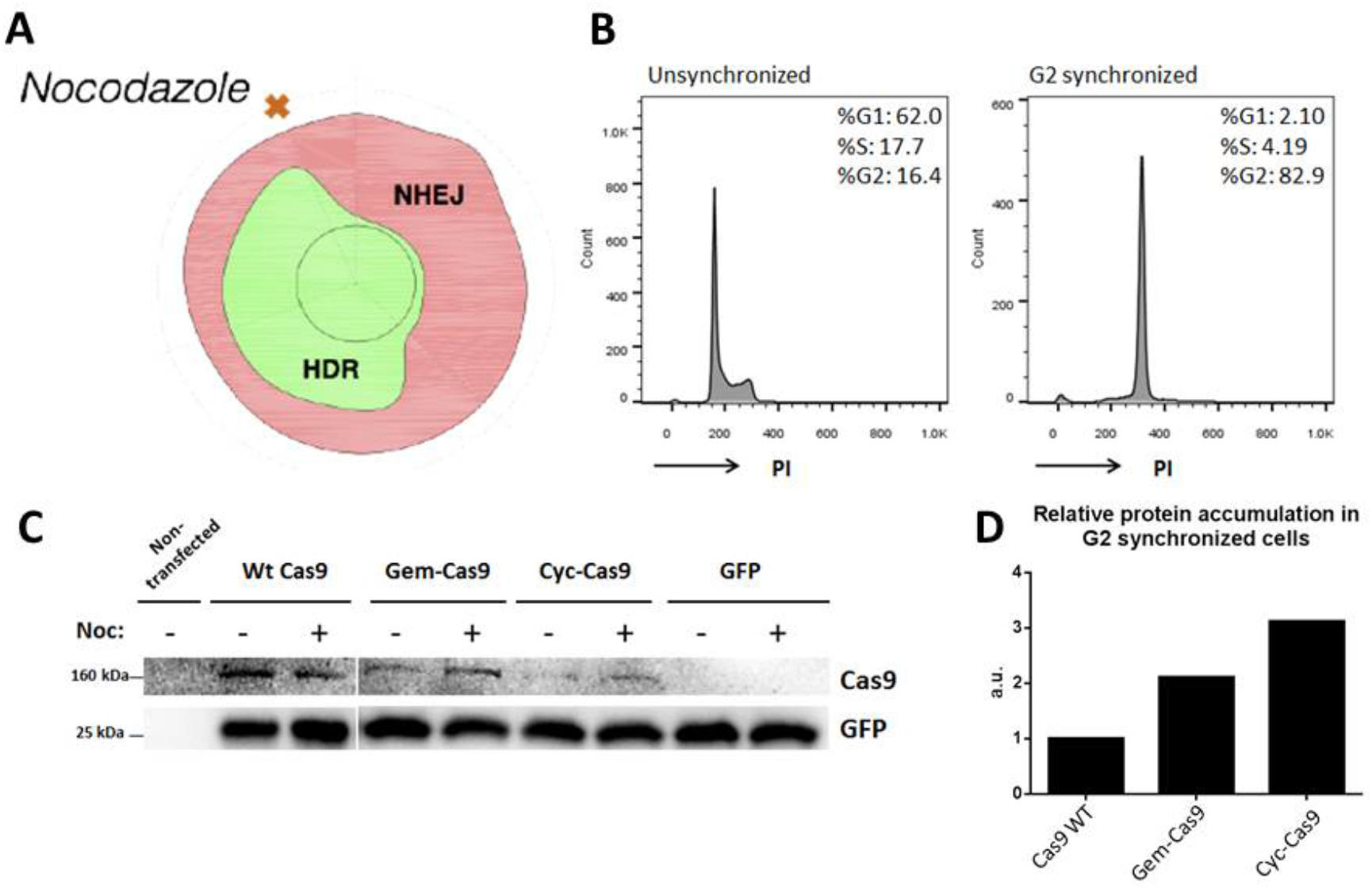
Analysis of protein degradation. (A) Scheme of cell-cycle arrest with nocodazole. (B) Cell cycle profiles of unsynchronized and G2-arrested cells, using a PI staining. (C) Western blot of protein extracts of unsynchronized and G2 arrested cells. GFP was used as a transfection and protein loading control. (D) Relative protein accumulation in G2-synchronized cells. Data are representative of 2 experiments.

## DISCUSSION

In this work, a strategy was presented to enhance the engagement of the HDR pathway on the repair of DSBs induced by CRISPR-Cas9. The goal was to design experimental tools to improve knock-in efficiencies, because the activity of HDR is relatively weak in actively dividing mammalian cells, when compared to other cell types or circumstances [30].

As mentioned before, different DSB-repair pathways are active at the same time. The competition between NHEJ and HDR for DSBs is influenced by several factors, including the cell cycle stage where the repair takes place [12, 28, 41 – 44]. The activation of the main repair proteins of HDR is controlled by cell-cycle-dependent mechanisms, whereas the NHEJ-related proteins are constantly active [10]. Thus, if Cas9 produces DSBs in the stages of the cell cycle where HDR is most active, an increase in HDR-mediated repair should be seen. Indeed, Lin *et al.* reported that the delivery of Cas9 in G2-arrested cells led to an increase in HDR-mediated repair [23]. The same goal is likely to be achieved in a simpler and less invasive way resorting to chimeric proteins composed of Cas9 and a motif known to promote the degradation of the protein in a cell-cycle stage-specific manner.

“Cell-cycle tags” were first used to monitor the cell cycle progression by fusing the Human Geminin’s 1-110 amino acids to induce the G1 degradation of GFP [38]. It was also applied to regulate the function of the uracyl-DNA glycosidase (UNG) in specific cell cycle stages, to address the issue of mutagenic repair of activation-induced cytidine deaminase (AID)-generated uracils. By fusing a UNG inhibitor to a Cyclin motif and another protein motif targeting the protein degradation in G1, Sharbeen *et al.* were able to compare UNG’s activity in specific stages of the cell cycle [45]. Additionally, after the planning of the fusions presented in this work, Gutschner *et al.* presented a Cas9-Geminin fusion that would be degraded in G1. This fusion has Cas9 fused at its C-terminus to the 1-110 amino acids of human Geminin and was shown to have a relative increase in G2-arrested cells, mimicking the Geminin’s degradation profile [46]. Another Cas9-Geminin fusion was presented, essentially with the same features, by Howden *et al.* [47]. Gutschner *et al.* started the evaluation of their Cas9-Geminin fusion by testing it on a classical knock-in experiment to correct a disrupted eGFP gene in the HEK293T cell line. The results show a maximum of 1.8-fold increase on HDR frequency. Next, the authors reasoned that the G2 synchronization would further increase knock-in efficiency and in an experiment designed to mutate a specific locus of the genome, they saw an approximate 4% increase on HDR rate, both for WT Cas9 and Cas9-Geminin. When compared to the WT Cas9, the fusion protein showed the same 1.8 fold HDR increase as in the experiment without synchronization.

In the present work, human Geminin was also chosen to induce a cell-cycle stage-specific degradation of Cas9, but in addition we tested the 2-87 amino acids of murine Cyclin B2, previously shown to induce G1 degradation [45]. It was of high importance to keep protein motifs as short as possible, to minimize the loss of function of Cas9. The Geminin-Cas9 fusion showed a statistically significant 40% increase compared to the WT Cas9. Although this result is similar to the one presented by Gutschner *et al.,* the modules fusion differ. First, the fusion protein published by Gutschner *et al.* has an extra N-terminus NLS besides the C-terminus NLS that is also present in the Geminin-Cas9 fusion presented in this work [25]. It remains to be shown if adding an extra NLS would boost the activity of the Geminin-Cas9 fusion protein here described; parenthetically, it should also be checked whether the Cas9-Geminin is inactive because of the design of the construct, which rendered the NLS internal and perhaps buried in the protein, despite the linkers used. Second, here HDR was measured using a cell line with a single copy of the HDR template inserted in the genome, whereas Gutschner *et al.* described a typical knock-in protocol using multiple transfected copies of an exogenous HDR template. The impact of the concentration of the HDR templates on the relative efficiency of the Geminin-Cas9 fusion compared to the WT Cas9 remains to be evaluated.

The other published Cas9-Geminin, by Howden *et al.,* lacked the addition of the extra C-terminus NLS and, compared to WT Cas9, showed similar levels of HDR frequencies. The main difference shown is the diminished NHEJ-mediated repair of induced DSBs, which is proposed as an advantage for knock-in experiments. Interestingly, Gutschner *et al.* show similar levels of indel formation by WT Cas9 and their Cas9-Geminin fusion. The decrease of NHEJ pathway activity on chimeric Cas9-induced DSBs was also shown here using the EJ5-GFP reporter cell line. The lack of NHEJ repair products may be explained not only by the timed cleavage activity of Geminin fusions in S/G2 and G2 that disfavors NHEJ repair, but also by the overall absolute cleavage activity: Geminin and Cyclin fusions are not present throughout most of the cell’s lifespan, giving rise to an overall drop in DSB formation.

The ratio between HDR and NHEJ provides an insight on the impact of the tested fusions in the recruitment of the DSB repair pathways. The NHEJ is three times more efficient than HDR in proliferating cells [48] and the goal is to shift the balance towards HDR. For both fusions tested here, Geminin and the newly reported Cyclin-Cas9, there is an increase on the HDR/NHEJ ratio, indicating that the overall balance between NHEJ and HDR mediated repair was shifted. Interestingly, the HDR/NHEJ ratio of the Cyclin-Cas9 fusion suggests that this fusion is overall less active, but has a higher specificity for HDR (Fig. 3D). This is probably due to the increased cell cycle-regulated degradation of this fusion compared to the Geminin Cas9 fusion, as shown by the ratio of protein degradation between unsynchronized and G2-synchronized cells (confirmed in two human-derived cell lines). Thus, the Cyclin-Cas9 fusion is a promising starting point for the optimization of a cell-cycle regulation of Cas9. Ongoing studies on the activity of Cas9 combining the post-translation regulation here described with cell cycle phase-specific transcription will soon be reported, including an evaluation of off-targets.

## ACKNOWLEDGMENTS

We thank Claúdia Andrade for data collection assistance and Jeremy Stark for sending us the reporter cell lines. This research was supported by the FCT PTDC/BEX-BCM/5900/2014 and IF/01721/2014/CP1252/CT0005 grants.

